# Agent-based modeling of nuclear chromosome ensemble identifies determinants of homolog pairing during meiosis

**DOI:** 10.1101/2023.08.09.552574

**Authors:** Ariana Chriss, G. Valentin Börner, Shawn D. Ryan

## Abstract

During meiosis, pairing of homologous chromosomes (homologs) ensures the formation of haploid gametes from diploid precursor cells, a prerequisite for sexual reproduction. Pairing during meiotic prophase I facilitates crossover recombination and homolog segregation during the ensuing reductional cell division. Mechanisms that ensure stable homolog alignment in the presence of an excess of non-homologous chromosomes have remained elusive, but rapid chromosome movements during prophase I appear to play a role in the process. Apart from homolog attraction, provided by early intermediates of homologous recombination, dissociation of non-homologous associations also appears to contribute to homolog pairing, as suggested by the detection of stable non-homologous chromosome associations in pairing-defective mutants. Here, we have developed an agent-based model for homolog pairing derived from the dynamics of a naturally occurring chromosome ensemble. The model simulates unidirectional chromosome movements, as well as collision dynamics determined by attractive and repulsive forces arising from close-range physical interactions. In addition to homolog attraction, chromosome number and size as well as movement velocity and repulsive forces are identified as key factors in the kinetics and efficiency of homologous pairing. Dissociation of interactions between non-homologous chromosomes may contribute to pairing by crowding homologs into a limited nuclear area thus creating preconditions for close-range homolog attraction. Predictions from the model are readily compared to experimental data from budding yeast, parameters can be adjusted to other cellular systems and predictions from the model can be tested via experimental manipulation of the relevant chromosomal features.

**Author summary:** Pairing of homologous chromosomes (homologs) is a key feature of multiple cellular processes including gene expression control, chromosome break repair, and chromosome segregation. Homolog pairing during meiosis is shared among all sexually reproducing eukaryotes. Mechanistic determinants of homology-specific chromosome alignment are presently unknown. We have developed an agent-based model where contributions of the entire chromosome set to the pairing process is taken into account, comprising both homologous and non-homologous chromosomal encounters. Incorporating natural chromosome lengths, the model accurately recapitulates efficiency and kinetics of homolog pairing observed for wild-type and mutant meiosis in budding yeast, and can be adapted to nuclear dimensions and chromosome sets of other organisms.

## Introduction

Double stranded DNA has an uncanny ability to find a homologous partner in DNA mixtures of staggering complexity. While homologs in somatic cells tend to occupy nuclear areas more distant than expected, somatic pairing nevertheless underlies important biological processes, including X chromosome inactivation and association of loci affected by genomic imprinting [1, 2]. During meiosis, homolog pairing is a key requisite for the separation of homologs to opposite spindle poles during meiosis I. When pairing is compromised, homologs fail to form crossovers, resulting in homolog nondisjunction and the formation of gametes with a surplus or deficit of one or several chromosomes. The resulting chromosomal imbalances are a leading cause for birth defects and still births [3].

In many organisms, meiotic pairing depends on recombination initiation via double strand breaks (DSBs), enzymatically induced by the spo11 transesterase [4]. DSBs typically occur at a multitude of chromosomal positions, at different positions in different cells. DSB processing is closely associated with the homology search, a process whereby 5’ resected DNA breaks assess homology between nearby chromosomes at the DNA sequence level [5]. If matched, DSBs are processed via homologous recombination into crossovers as well as other recombination products [6]. Crossovers involve the reciprocal exchange of chromosome arms between homologs at allelic positions [7]. In addition to providing physical linkage between homologs and ensuring their attachment to opposite spindle poles, crossovers also increase genetic diversity [6].

The timing and genetic requirements of homolog pairing have been extensively studied in several organisms [6–10]. In budding yeast, homologs are somatically paired in G1-arrested cells, unpair during premeiotic DNA replication and commence re-pairing as cells initiate homologous recombination [10–12]. Around the time when homolog pairing is established, chromosomes also undergo rapid movements throughout prophase I, as a prerequisite for efficient pairing. Movements of chromosomes during meiotic prophase I are mediated by motile cytoplasmic filaments, actin in budding yeast and the dynein-microtubule complex in most other organisms, which drag chromosome ends (telomeres) through the semi-fluid nuclear envelope.

Cytoplasmic motorproteins mediate nuclear chromosome movements due to the attachment of chromosome ends via the conserved SUN-KASH protein complex where SUN proteins interact with chromosome ends and reach across the inner nuclear envelope whereas KASH proteins span the outer nuclear envelope providing a link between SUN proteins and the cytoplasmic filaments [13–15]. Pairing is completed around the time when both DSB ends have undergone strand exchange, giving rise to double Holliday junctions, a critical precursor of crossovers [10, 12]. Both recombination initiation and DSB processing into joint molecules including single end invasions and double Holliday junctions is required for homolog pairing, even though crossover formation itself appears to be dispensable [8, 10, 12, 16, 17].

The synaptonemal complex (SC) is a proteinaceous structure that assembles between paired homologs and juxtaposes their axes closely at 100 nm, a distance conserved in most taxa [6, 8–10]. Homologous chromosomes are considered as paired once they have associated at a distance less than or equal to 400 nm, corresponding to the distance between co-aligned homolog axes in absence of the SC [8].

There are many challenges to experimentally monitor homolog pairing. Fixated, surface-spread cells exhibit superior resolution, but they do not provide insights about chromosome dynamics during the pairing process and nuclear architecture may become distorted during sample preparation [12, 18]. Tracking of chromosome trajectories in live cells is hampered by limited resolution and potential effects of phototoxicity [19]. Importantly, such studies are limited to a small subset of chromosomes due to the necessity to fluorescently label individual chromosomes [10, 16, 20].

Little is known about the molecular mechanism(s) of homolog pairing. Several mathematical models have been put forward examining potential contributions of molecular processes to homolog pairing, including telomere attachment to the nuclear envelope, chromosome bending stiffness or polymer chains exhibiting an excluded volume repulsive potential [19, 21–24]. Moreover, a cellular automaton model was developed that examines random searching via chromosome shuffling [25]. Importantly, existing pairing models cannot be validated due to inaccessibility of individual pairing events to experimental analysis.

Here, we have developed an agent-based model (ABM), in which movements and interactions initiated by individual chromosomes (i.e. agents) are simulated, thereby recapitulating the interaction dynamics of an entire nuclear chromosome ensemble. Rather than making assumptions about pairing dynamics along individual chromosomes, we use differential equations based on first principles that govern the movements of chromosomes as an outcome of interactions with all other chromosomes within the same nucleus. Our model allows for the analysis of individual trajectories for all chromosomes throughout meiotic prophase I, facilitating comparison with experimental data. Modifications of various chromosomal parameters, including chromosome number, size, and movement velocity as well as attractive and repulsive forces provide insights into the contributions of each of these factors.

## Materials and methods

### Modeling Approach

Our homologous pairing model considers three contributing processes, i.e., chromosome interactions (homologous and non-homologous collision dynamics), chromosome translation (self-propelled, directed movements), and random chromosome motion (thermal noise within the nucleus) (Fig. 1). By deriving a model entirely from these first principles, we avoid the introduction of mathematical parameters not apparent from the underlying biological process. A key feature of our model is the inclusion of attractive and repulsive forces representing processes that stabilize homologous pairing or disrupt non-homologous interactions, respectively. Rather than focusing on a single homolog pair, our model captures trajectories of complete chromosome sets throughout prophase of meiosis I, facilitating comparison with experimental data and adjustments to species-specific features such as chromosome size and number as well as nuclear dimensions.

**Fig 1.**
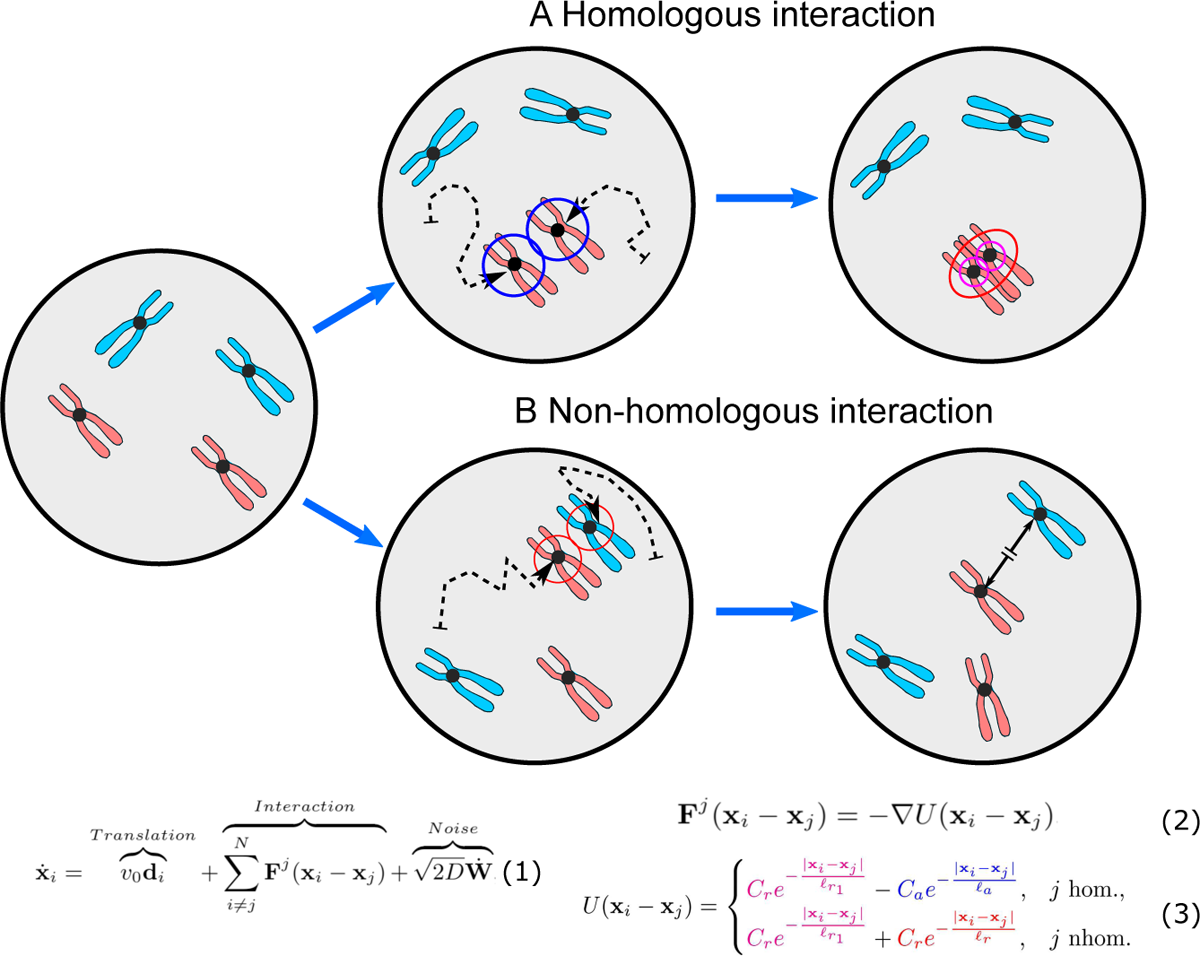
Determinants of chromosome dynamics during the meiotic homology search. All chromosomes move about the cell nucleus and while searching for their homologous pair perform a continuous random walk (dashed arrows) within the confines of the nuclear envelope (bold circles), with velocity and changes in direction determined by interactions between chromosomes as well as thermal noise. (A) When homologs enter each other’s attractive radii (i.e., centers are 400 nm apart), they exert attractive forces (Eq. (3); j hom) on each other which move them closer until they reach their respective exclusive radii (magenta; here 50 nm) keeping them at a constant distance. They subsequently continue their effective “random walk” in a paired status moving as a single non-homologous chromosome pair with respect to all other chromosomes in the nucleus. (B) When a chromosome enters the repulsive radius of a non-homolog (red; here 400 nm), the repulsive force in (Eq. (3); j nhom) diverts their movement at the angle of their encounter with new velocity proportional to the minimum interaction distance. For illustration, the ring colors match the terms in the Morse potential governing the interactions between chromosomes in Eq. (1)-(3); also see Fig 2.

To compute the net motion of meiotic chromosome sets, we consider only pairwise interactions between chromosomes at short distances as provided by a semi-dilute solution [26, 27]. A two-dimensional model is developed to match the available two-dimensional experimental measurements on fixated, surface-spread yeast nuclei used to calibrate the model [12]. For the dynamics of the center of mass, x*_i_* R^2^, a coupled system of *N* equations can be proposed as

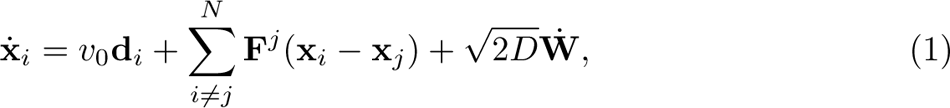

where *N* gives the number of chromosomes and F*^j^* is the force between two chromosomes, which is obtained as the negative gradient of the potential energy *U*

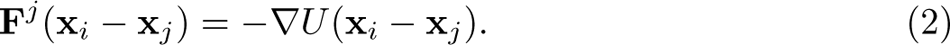

The potential energy is defined using the Morse potential [28]

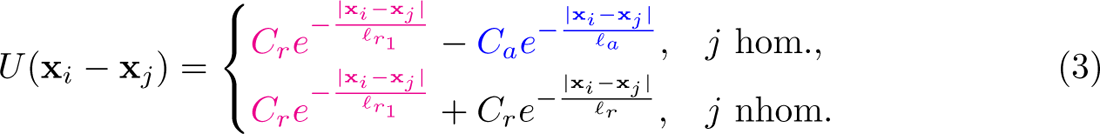

Three components in Eqs. (1) are noteworthy. First, there is a translational velocity term (*v*_0_d*_i_*) defining a chromosome’s present straight-line motion as one component of its total velocity. We refer to this as the chromosome’s translational velocity because it is straight-line motion in the direction the chromosome is already moving. The velocity (*v*_0_) is multiplied by a chromosome’s dimensionless orientation d*_i_* = v*_i_/*lv*_i_*l, a unit vector in the direction of the chromosome’s velocity. d*_i_* can be changed by collisions with other chromosomes or with the nuclear envelope as well as by thermal noise. In living cells, chromosome movement is typically generated by cytoplasmic motile filaments which are connected to chromosome ends (telomeres) via a protein complex that traverses the semi-fluid nuclear envelope (see Introduction) [6, 23, 29, 30]. In our model, chromosome end attachment to the nuclear envelope is reflected by the two-dimensional features of the nuclear volume simulating movement along an invariable *z*-surface.

Second, interaction forces between chromosomes are captured by Eqs. (2) and (3). The force F is derived from the potential energy defined by the Morse potential *U* and is singularly determined by the distance between homologous or non-homologous pairs of chromosomes x*_i_* x*_j_* [28]. A Morse potential is defined as a difference of Yukawa potential energies, assigning to homologous chromosome pairs a weak attractive (i.e. associative) force strength *C_a_* (blue ring in Fig. 1A; here *ℓ_a_* = 400 nm) and a weak repulsive (i.e. excluded volume) force strength *C_r_*_1_ acting at very short distances and ensuring that paired homologs are kept at a fixed center-to-center distance rather than overlapping (purple ring in Fig. 1; here *ℓ_r_*_1_ = 50 nm). Using a purely repulsive force derived from an energy potential to enforce an excluded volume constraint was also successfully used recently to study chromosome pairing in [24]. Accordingly, interactions between homologs are determined by their distance: namely, attractive with increasing strength as distance decreases from 400 to 50 nm, and repulsive at or below 50 nm. For non-homologous chromosomes, the potential energy is provided by a purely repulsive (i.e. dissociative) force strength *C_r_* that may model an abortion of strand exchange over short distances (red ring, in Fig. 1B; here *ℓ_r_* = 400 nm) as well as the excluded volume constraint *C_r_*_1_ acting at a distance of *ℓ_r_*_1_ = 50 nm (see Table 1). Following a non-homologous encounter, chromosomes move away from each other while maintaining their orientation with the new velocity proportional to the interaction distance. Mathematically, the Yukawa potential has a simple formula for its derivative defined in Eq. (2) which gives the corresponding interaction force and allows for straightforward numerical computation.

**Table 1.** Table of parameters used in the simulations. Movement speed for *spo11* hypomorph from [14].

| Parameter | Dim. Value | Nondim. Value |  |  |
| --- | --- | --- | --- | --- |
|  |  | Base | *spo11* (Fig. 10B) | *spo11*(Fig. 10C) |
| Nucleus (domain) radius $L$ | 3.25 $\mu\text{m}$ [12] | 3.25 | 3.25 | 3.25 |
| Characteristic length scale $L_e$ | 1 $\mu\text{m}$ [15] | 1 | 1 | 1 |
| Diffusion constant $D$ | $10^2 \text{ nm}^2/\text{s}$ [51] | 0.0001 | 0.0001 | 0.0001 |
| Movement speed $v_0$ | 300 nm/s [29] | 0.3 | 0.3 | 0.231 |
| Length scale, attractive force $\ell_a$ | 400 nm [8] | 0.4 | 0.4 | 0.4 |
| Length scale, repulsive force (non-hom. chromosomes) $\ell_r$ | 400 nm [8] | 0.4 | 0.4 | 0.4 |
| Length scale for repulsive force (hom. chromosomes) $\ell_{r_1}$ | 50 nm [6] | 0.05 | 0.05 | 0.05 |
| Attractive Strength $C_a$ | N/A | 0.005 | 0.0017 | 0.0017 |
| Repulsive Strength for non-hom. chromosomes $C_r$ | N/A | 0.005 | 0.0017 | 0.005 |
| Excl. Vol. Strength for all chromosomes $C_{r_1}$ | N/A | 0.05 | 0.05 | 0.015 |
| Boundary repulsion strength $C_b$ | N/A | 0.15 | 0.15 | 0.15 |
| Boundary repulsion length $\ell_b$ | N/A | 0.05 | 0.05 | 0.05 |

The form of the interactions as a Yukawa decaying exponential ensures that forces between chromosomes become rapidly negligible beyond the effective interaction radius *ℓ_a_* or *ℓ_r_* respectively. Each of the three forces are at their respective maxima when chromosomal centers of mass overlap, and decrease exponentially from there, falling, e.g., to ~ 1*/*3 of their maxima when their distance has reached the effective radius. Beyond this effective radius they are essentially negligible compared to chromosome motion because these components of velocity are significantly smaller than the translation velocity, *v*_0_, or the random motion term. For example, when chromosome centers of mass are separated by three effective length radii, forces decrease by an order of magnitude from their maxima. Thus, all computed forces represent short-range interactions that do not affect the motion of either homologous or non-homologous chromosomes beyond one effective length scale, as further evident from Supplemental Movies S1 to S3 of representative simulation runs (see Fig. 2; below).

**Fig 2.**
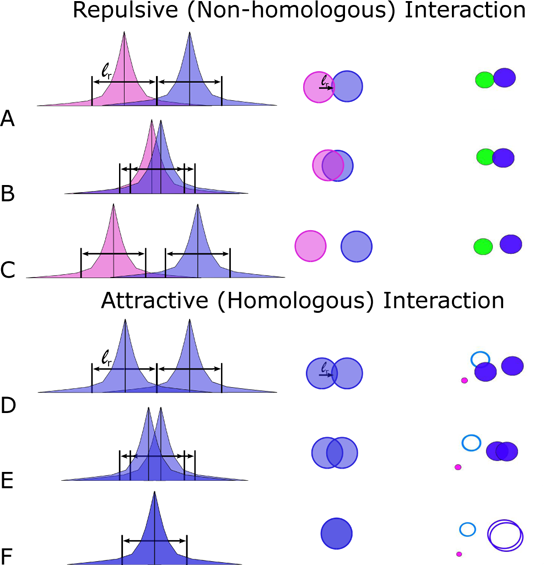
Strength of exponentially decaying forces between non-homologous (short-range repulsive) and homologous (short-range attractive) chromosomes. The vertical black line represents the center of mass, x*_i_*, and the colored regions indicate each chromosome’s force modeled as a decaying exponential. The first column shows the strength of the forces. The second column shows a cartoon illustration of the corresponding chromosome dynamics when they meet, and the third column is a still image from Supplemental Movie S1 Video. The solid circles on the righthand side indicate the effective interaction radius (see Table 1) beyond which the force is negligible. (A-C) Illustration of exponentially decaying forces when two non-homologous chromosomes meet. The repulsive force felt by a participating non-homologous chromosomes corresponds to where the center of mass crosses into the other chromosome’s repulsive region. (D-F) Similar illustrations but for an attractive homologous interaction.

Within living cells, attractive and repulsive forces may result from a combination of diverse interactions. These interactions primarily entail homology search and strand exchange between a resected double-strand break (DSB) and intact template DNA. However, they can also encompass interactions between fully intact double-stranded DNA molecules. Attractive forces manifest during the formation and elongation of a heteroduplex between a DSB and a homologous template. This process is initiated by microhomologies, typically consisting of 8 base pairs [31]. On the other hand, repulsive forces involve the dissociation of heteroduplexes containing internal or flanking mismatches, a phenomenon facilitated by ATP hydrolysis [32]. Factors influencing the success of pairing and strand exchange include the degree of coiling in both the template and invading DNA [33] and, at closer proximity, electrostatic interactions [34]. Proteins, such as histones [35], RecA orthologs [32] and mediator proteins [36], diverse ATPases [37], and the mismatch repair machinery ([38, 39]), can modulate both attraction and dissociation processes.

Third, there is a term in Eq. (1) accounting for th_√_e<u>rm</u>al noise within the nucleus. This is a small white-noise process where W° ∼ *N* (0*, dt*) can be modeled as a normally distributed random variable with mean zero and variance proportional to the time step. Though the chromosomes are tethered to the nuclear envelope, their motion is not linear and thermal noise plays an active role at this small scale. This is best modeled as a random walk. Although the net motion is not that of a random walk, this component contributes biologically more relevant overall motion. This formulation has been successfully used to capture thermal noise in other models describing microscale biological processes [40–46].

Our approach replaces microscopic details of the physical chromosome shape with a representation of the dynamics of its center of mass, which resembles the green fluorescent protein (GFP) dot in the experimental setting. This allows for chromosome movement to be the direct response of interaction frequencies between homologous and non-homologous chromosomes throughout the nucleus. Our model is simplifying the nucleus from three to two dimensions, providing a direct correspondence to the two-dimensional nature of the experimental data set used to calibrate the model where biological sample preparation involved fixation and flattening of the nucleus [12]. Furthermore, our modeling framework treats homolog pairing as an endpoint and does not consider the ensuing homolog segregation during meiosis I, again facilitating comparison to experimental data where cells were arrested at the pairing stage due to the absence of the meiotic progression factor Ndt80 [12, 16].

### Parameter Choice

Table 1 displays the parameter values used in the initial simulations. While our model is adaptable to a wide range of parameters, it is important to identify realistic values for comparison with specific experimental data. Meiosis in the budding yeast *S. cerevisiae* (the organism which provided the experimental data set used for model calibration) involves 16 homolog pairs or 32 individual chromosomes, corresponding to 32 coupled equations, each simulating the dynamics of the center of mass for an individual chromosome. Chromosomes may come in close contact with the nuclear boundary without ever crossing it due to an appropriate value set for the boundary repulsion strength and characteristic length scale using a purely repulsive Morse potential (similar to pure repulsion case in Eq. 3). While the intact yeast nucleus is roughly 2*µ*m in diameter, it is set here to 6.5*µ*m to account for its flattening and spreading during experimental data collection [5, 12] facilitating comparison with recent experimental data to establish model validity. Flat nuclei in the simulation result in chromosome motion exclusively within the two-dimensional cross-section shown in Figure 1. Experimentally determined distances between GFP-tagged chromosomes range between 200 nm (the resolution limit of light microscopy) and 6.0*µ*m [12]. Chromosomes are tethered to the nuclear envelope, and the simulation effectively tracks the chromosome movement in this two-dimensional cross-section.

While attractive and repulsive interactions between homologous and non-homologous chromosomes, respectively, are presently not accessible to direct measurements, their length scales can be estimated from features that have been determined in several biological systems (above). Many of these interactions involve evolutionarily conserved components of the homologous recombination machinery for which measurements are available [6]. We use a “symmetric” model where attractive and repulsive forces act at the same length so as not to overvalue one type of interaction. The attractive length radius (*ℓ_a_*) is set to 400 nm corresponding to approximately 800 nucleotides of fully extended, single stranded DNA (assuming 0.50 nm/bp; [47, 48]). During the homology search, such 5’ resected DSB tentacles may reach out from the broken chromosome to other chromosomes to probe homology [9]. The exponential decay of both attractive and repulsive forces further reflects ranges of single stranded DNA at meiotic DSBs ranging from 500-1500 nucleotides [48, 49] Thus, two chromosomes with centers of mass within 400 nm will initiate the homology search. The minimum homology length requirement for strand capture entails 8 nucleotides of homology; such initial contacts are thought to become extended in nucleotide triplet steps [31]. Therefore, a relatively short DSB tip region appears to be sufficient to make the first contact followed by homolog movement toward one another by heteroduplex extension. Once homologs are paired, they remain so henceforth and assume a single equation of motion ensuring they do not break apart, with the very short distance repulsive force keeping them at a constant distance of 50 nm which approximately corresponds to the conserved 100 nm width of the synaptonemal complex [6].

Likewise, the repulsive length (*ℓ_r_*) is set to a 400 nm center of mass radius under consideration that invading resected DSB tentacles also generate a dissociative (repulsive) force. Dissociation of strand exchange due to insufficient homology likely involves the same forces as attraction, successful strand exchange, though resulting in a net movement in the opposite direction. Thus, 400 nm is chosen as the distance needed to determine if an approaching chromosome is non-homologous. The repulsive force further provides a parameter that minimizes the time spent by non-homologous partners in close vicinity. We note that both attractive as well as repulsive forces are significant only over short distances, i.e., when chromosome centers of mass are less than 400 nm apart, which corresponds to less than 1/15 of the maximum distance provided by the 6.5*µ*m diameter of the nuclear area (Fig. 3).

**Fig 3.**
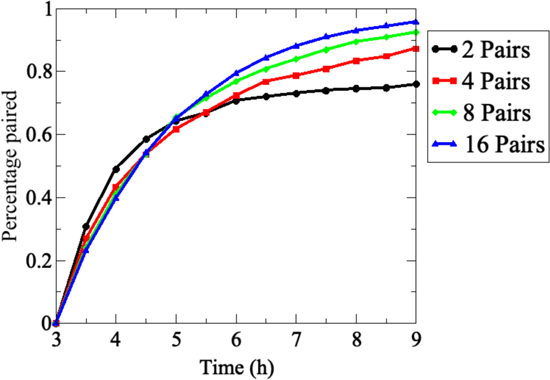
Effects of chromosome number on pairing efficiencies and kinetics. Pairing frequencies for the indicated number of homolog pairs in 200 realizations are combined to compute the percent of homologs paired at the indicated time points. Average pairing levels of the indicated number of equally sized homolog pairs. Error bars are omitted for clarity.

Our choice for the strengths of homologous and non-homologous interaction forces is further motivated by kinetic considerations. Non-homologous interaction strength is directly related to the time spent exploring that interaction by a given chromosome. Thus, doubling the repulsive strength cuts the time spent in proximity in half for non-homologous chromosomes (see Fig 2). By contrast, interactions between homologous chromosomes requires a longer time period for the assessment of homology. Parameters also need to reflect features of the decision process whether or not two chromosomes are homologs (e.g., parameters should not prevent prolonged association of non-homologous chromosomes), but also not result in chromosomes jumping apart. It is further noteworthy that a given chromosome has an attractive interaction with one homologous chromosome, but a repulsive interaction with up to 30 non-homologous chromosomes, even though in the semi-dilute scenario, interactions are assumed pairwise meaning that they typically occur with only one other chromosome at a time.

Apart from collisions, the homology search involves the following processes. Directed, chromosome movements increase the probability of chromosomal encounters allowing them to probe homologous and non-homologous DNA partners. The velocity of chromosome movements at 300 nm/s is taken from live-cell imaging of meiotic chromosome movements which range between 200 and 500 nm/s [14, 15, 29]. Importantly, prophase I chromosome movements enhance chromosomal encounters, whereas the homology search for a given chromosome with 400 nm attractive radius would limit to approximately 5% of the nuclear area without movements [50]. The parameter for chromosome motion due to thermal noise is provided from the diffusion constant of interphase chromatin which ranges between 10^2^ and 10^3^ nm^2^/sec [51].

At the start of each simulation, chromosomes are assigned unit velocities and random initial placements throughout the nucleus using a basic exclusion algorithm where a chromosome is placed and then its center of mass distance to all other chromosomes previously placed is computed [21]. If any initial distance is below 400 nm indicating pairing, another random location is chosen for the respective chromosome to avoid overlap in the initial chromosome placement, repeating this process until all chromosomes have been placed. Notably, the model is easily adaptable to alternative initial placement exclusion distances.

## Results

### Initial model and comparison with experimental data

In the reference experiment, homologous and non-homologous chromosome interactions, respectively, were monitored by retrieving cell aliquots from semi-synchronous meiotic cultures which either carry two GFP-tagged copies of chromosome III or single copies of GFP-tagged chromosomes III and II [12]. Pairing was inferred from the frequency of cells with GFP dots separated by less than 400 nm in the strain carrying GFP-tags in homologs adjusted for fortuitous co-localization as derived from measurements in cultures carrying GFP-tags in the non-homologous chromosomes pairs. This analysis suggested that homolog pairing is at a minimum around *t* = 3*h* after transfer to meiosis medium, corresponding to the time of pre-meiotic S-phase, and reaches maximum levels at *t* = 7.5*h*, when most cells have entered the pachytene stage [12].

To model the complete homology search process throughout prophase I, we simulated chromosomal dynamics using Eq. (1)-Eq. (3). The numerical approach was designed in MATLAB [52]. A base code for the wild-type dynamics is available at [53]. Importantly, the simulation allows concurrent tracking of all 16 homolog pairs and comparing their pairing dynamics to appropriately matched non-homologous partners within the same nucleus. Running Monte Carlo simulations with up to *N* = 32 chromosomes, typically 200 realizations were considered and averaged. We began assessing the validity of our model under the simplification that all chromosomes are of uniform length corresponding to the measured length of a mid-sized yeast chromosome at the pachytene stage [15] and a uniform reach corresponding to 400 nm. Parameters in Table 1 are chosen for the best correspondence with experimental data following exploration of the parameter space.

We initially modeled pairing with a minimum set of two pairs of homologous chromosomes and subsequently increased the number of chromosome pairs to 16. Homologs were considered as paired when they became stably juxtaposed at 400 nm (above). Exploring pairing dynamics in dilute (less then 4 pairs) to semi-dilute (4 or more pairs) conditions. This study revealed unexpected effects of chromosome numbers on the efficiency and kinetics of pairing.

The model was started at the *t* = 3*h* time point, corresponding to minimum pairing levels in the experimental data set. For the scenario with 2 homologs, only 75% of homolog pairs have completed pairing by *t* = 7*h*, and increases at later times are negligible. An increase of homolog pairs to 4 increased the efficiency of pairing somewhat and this increased further in the presence of 8 or 16 homologs pairs per nucleus resulting in a substantial increase in maximum pairing levels. This reveals the qualitative difference between the dilute (2 pairs) and semi-dilute regime (4 or more pairs) in chromosome dynamics. Accordingly, with at least 8 homolog pairs, essentially all homologs have completed pairing by *t* = 9*h*. Thus, in the dilute scenario, isolated movements without chromosome interactions are predominant, whereas in the semi-dilute scenario chromosome interactions dominate motion and such interactions are a prerequisite for successful homolog pairing.

Having established that chromosome number affects both pairing efficiency, we next explored additional features of the scenario with 16 homolog pairs, i.e. the chromosome set in diploid budding yeast. In this analysis, homolog pairs were arranged based on their initial distances in ascending order. Homolog distances were then recorded over time and plotted at four time points corresponding to those examined experimentally for chromosome III (Fig. 4A). Each data point represents average distances from 200 realizations. Initial distances between homologs in the simulation range from 0.7*µ*m to ∼ 6*µ*m (Fig. 4A). Modeling suggests that chromosomes initially separated by 0.7*µ*m to 4*µ*m (chromosome indices 1 to 12) have completed pairing by *t* = 5*h*, within 2 hours, whereas chromosomes separated by a larger distance require up to 4 hours to complete pairing (Fig. 4A). These results suggest a nonlinear relationship between initial homolog distance and pairing kinetics. Moreover, there appears to be a critical initial distance threshold above which homolog pairing becomes very slow.

**Fig 4.**
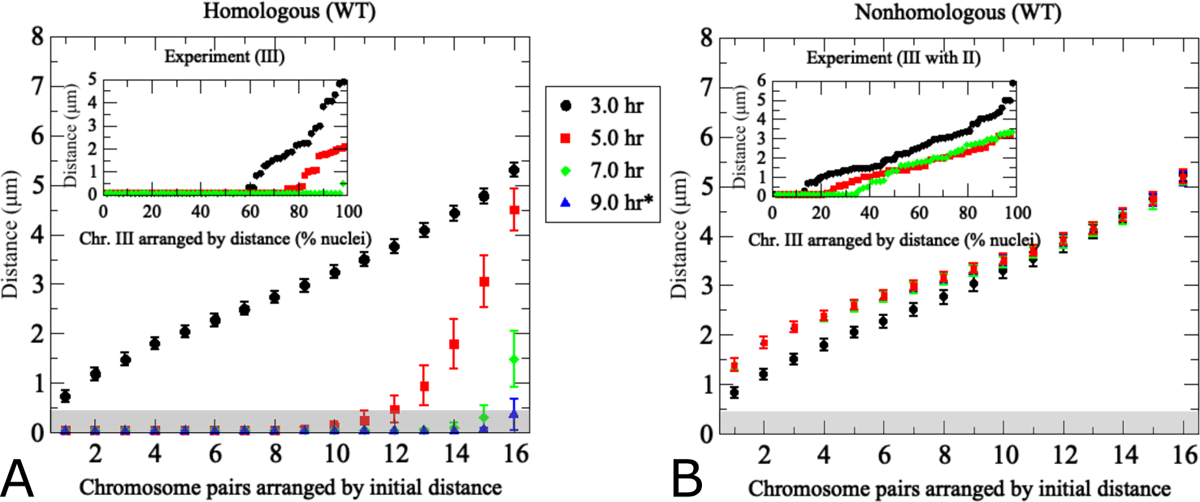
The pairing model captures experimentally determined association kinetics between homologous and non-homologous chromosomes during wild-type meiosis. (A) Modeling of pairing between 16 uniformly-sized partner chromosomes. For each modeling run, chromosome distances at *t* = 3*h* are used to uniquely index homologous chromosome pairs, which are arranged based on ascending initial distances along the *x*-axis. The *y*-axis indicates average distances from 200 realizations of the agent-based model Eq. (1) – Eq. (3) at the time points indicated by the color code. Error bars indicate standard deviations. The inset shows experimentally determined distances between a single pair of GFP-labeled yeast chromosome III measured in a synchronized meiotic culture in fixated nuclei (*n >* 100) (data from [12]). Nuclei are arranged based on distances between homologous GFP signals at a given time point. Note that x-axes are different in the inset due to the fact that in the experiment cells were observed at a given time point and then discarded, whereas in the simulation the same cell was tracked over time. (B) Results from modeling of distances between non-homologous chromosome pairs in the same nuclei analyzed in (A). For non-homologous pairs, each homolog partner is matched with a non-homologous chromosome that at *t* = 3*h* exhibits a distance optimally matched to that with its cognate homolog partner. Inset (B) shows experimentally determined distances between non-homologous GFP-tagged budding yeast chromosomes II and III (error bars indicate SD). For details on experimental conditions see [12]. The pairing distance is highlighted in gray at 400 nm. *Note the experimental data do not include information for *t* = 9*h*.

Distances between non-homologous chromosome partners were also monitored in the same simulation. Data points in this case represent any non-homologous chromosome initially placed at the same distance as the homologous partner chromosome in the same nucleus (Fig. 4B). Accordingly, initial distances between non-homologous and homologous pairs are similar, yet non-homologous distances remain unchanged over time whereas homologous distances become progressively smaller. This confirms that the simulation is indeed specific for homologous pairing.

Comparison between simulation and experiment indicates good qualitative and quantitative convergence for homologous and non-homologous association kinetics (see insets Fig 4). Importantly, non-homologous GFP signals remain at similar distances throughout meiosis in both experiment and simulation, with little changes in inter-chromosomal distances. We conclude that the modeling framework reasonably recapitulates the chromosome dynamics associated with homologous pairing throughout prophase I. Notably, however, modeled pairing kinetics are derived from the entire set of 16 homolog pairs within the same nucleus whereas experimental data represent distances between a single GFP-tagged chromosome with its homologous partner or a single representative non-homologous partner measured in different nuclei and from different cultures.

Several additional differences between experimental data and modeling are noteworthy. In the simulation, all chromosome pairs are placed to an unpaired starting position and released into the moving chromosome ensemble. The experiment, by nature represents a more complex situation: (i) Due to the incomplete synchrony within a meiotic culture, a cell sample taken at a given time point comprises a mixture of cells that have progressed to various degrees in the pairing process. (ii) Moreover, prior to entry into meiosis, homologs are paired somatically in G1-arrested cells, potentially as a special case of yeast meiosis [10–12], meaning that both laggards that have not yet completed unpairing and those pairing ahead of the population average will artificially increase the pairing frequency. Accordingly, a substantial subset of cells exhibits homologous pairing even at the time of minimum pairing levels, a substantial subset of cells exhibits homologous pairing (see insets of Fig. 4A), either because they are still somatically paired or because they have already progressed to post-replicative pairing. (iii) While the model places unpaired chromosomes randomly on an idealized two-dimensional nuclear area, the experimental “area” is created through spreading and/or squashing a three-dimensional cells in an imperfectly controlled manner potentially compressing homologs that are separated along the *z*-axes, a complication experimentally addressed via comparison with cells harboring GFP signals on non-homologous chromosomes. Thus, unpaired homolog pairs may end up in close vicinity, even though this close vicinity may not have existed when the nucleus was in its original 3D state. This experimental artifact is further evident from substantial frequencies of pairing between non-homologous chromosomes in the experiment (see inset Fig. 4B).

### Modeling predicts delayed pairing for the three smallest yeast chromosomes

The 16 budding yeast chromosomes range in length from 250 kilobasepairs (kbp) to 1,500 kbp. Chromosome size affects several meiotic processes, including the timing and/or density of initiating DSBs and crossovers [54, 55]. Chromosome length may further affect the cumulative range of attractive and repulsive forces as well as the available space for other chromosomes to move in [56]. We therefore took the length of yeast chromosomes into account by scaling all attractive and repulsive radii in Table 1 by an individual chromosome’s relative length as compared to the average chromosome length. The larger area takes into account the higher number of DSBs engaged in the homology search along a larger chromosome, even though the reach of an individual DSB would likely remain unchanged. The inner repulsive radius enforcing binding distance between homologs were kept at 50 nm, independent of chromosome length, consistent with the uniform width of the synaptonemal complex.

As before, distances between homologous chromosomes were plotted as a function of time, but results were sorted by increasing chromosome lengths rather than by initial chromosome distance (Fig. 5A). For one of the realizations of the entire homology search of size-adjusted homolog pairs, see Supplementary Movie S1 (https://zenodo.org/records/10246589). Again, initial chromosome distances are on average 3*µ*m for all chromosomes, independent of chromosome length, corresponding to half of the diameter of the cell nucleus. As time progresses, chromosomes longer than 400 kbp have completed homologous pairing by *t* = 7*h* in essentially all nuclei, whereas the three shortest homolog pairs which range between 250 and 320 kbp remain unpaired, even at *t* = 9*h*, as suggested by their final distances of *>* 400 nm (Fig. 5). Thus, the three shortest chromosomes take substantially longer to pair than the rest of chromosomes, revealing a nonlinear effect of chromosome length on pairing kinetics. Importantly, these distinct kinetics are limited to small homolog pairs, but are absent for the same chromosomes and their equidistant non-homologous chromosomes in the same nuclei (Fig. 5B). Hence, our model predicts that smaller chromosomes exhibit unique pairing properties.

**Fig 5.**
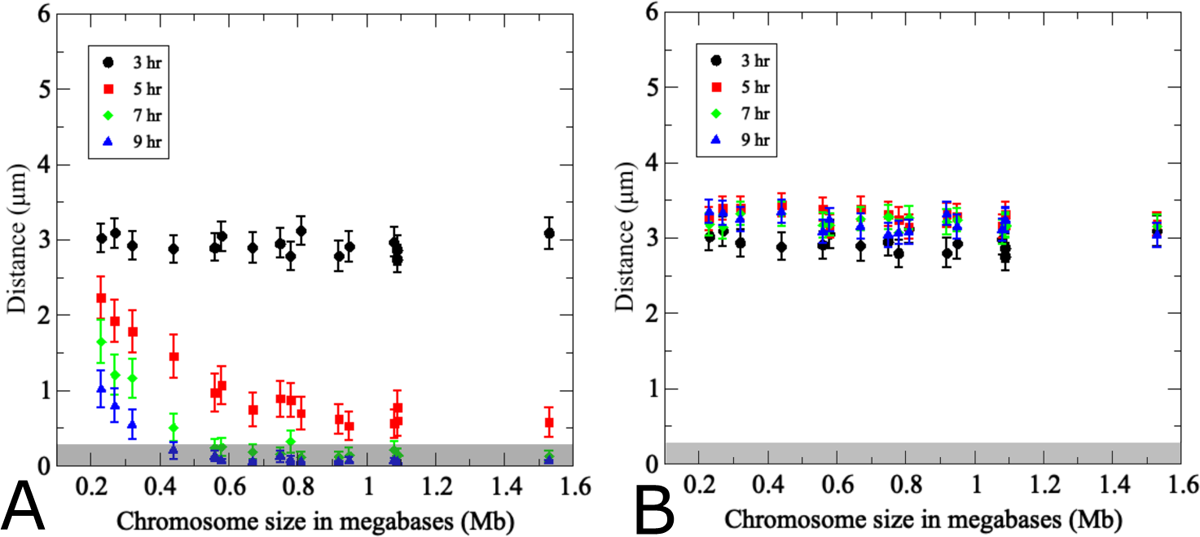
Effects of chromosome size on pairing kinetics and efficiency. (A) Average distances over time between homolog pairs where repulsive and attractive radii are adjusted proportionally to the sizes of actual yeast chromosomes. Note that the three smallest homolog pairs are markedly slower in achieving pairing than all other chromosomes. (B) Non-homologous chromosomes equidistant to each of the two homologs were identified at *t* = 3*h* and their distances were monitored throughout the simulation in the same set of model nuclei monitored in (a). (200 realizations, error bars indicate SD). The gray rectangle highlights pairing distances at or below 400 nm.

Several additional features of the model are further evident from inspection of the Supplemental Movie S1 Video: Both attractive and repulsive forces affect the path of other chromosomes only at short distances, as the path of a given chromosome is only altered when their effective radii overlap. Moreover, even when homologous chromosomes approach each other, they may remain closely aligned for extended periods, but then fail to complete pairing at that time and separate again, e.g., due to the force of a repulsive interaction with a non-homologous chromosome. Such dynamics may explain the mixed association previously observed during experimental live-cell imaging studies where GFP-tagged homologs approached and separated without completing pairing [19].

### Effect of chromosome movement velocity on pairing

We next used our model to examine the role of chromosome translation velocity, *v*_0_, on pairing efficiencies and kinetics. Even though faster chromosome movements would be expected to uniformly accelerate homolog pairing, these simulations identified a velocity threshold for pairing, which particularly impacts larger chromosomes. In the simulations described previously, the velocity parameter in Eq. (1) was set to *v*_0_ = 300 nm/s (see Table 1). Reducing the chromosome translation velocity by 50% (to 150 nm/s) essentially eliminates pairing when average pairing frequencies for size-adjusted chromosomes were monitored over time (red line; Fig. 6A). Increasing chromosome velocity in 30 nm/s increments to 210 nm/s results only in minor increases in pairing efficiencies, with on average only 30% of homolog pairs achieving pairing by *t* = 9*h* (Fig.6A), blue). Surprisingly, the four smallest chromosomes (sized below 500 kbp) disproportionately contribute to pairing at velocities at or below 210 nm/s, whereas pairing is essentially abrogated for mid-sized and larger chromosomes (Fig. 6B). When chromosome translation velocity is further increased by 30 nm/s to 240 nm/s, this results in a dramatic improvement of pairing efficiency for all chromosomes, with now ~ 50% of homolog pairing completed by *t* = 9*h*, regardless of chromosome size (orange; Fig. 6A,B). Pairing efficiency and kinetics can be further improved for all chromosomes by an increase of velocity to 270 nm/s (light blue), whereas neither a further 30 nm/s incremental increase nor a doubling in chromosome velocity to 600 nm/s has substantial effects on pairing kinetics or efficiency (Fig. 6A,B; black and purple).

**Fig 6.**
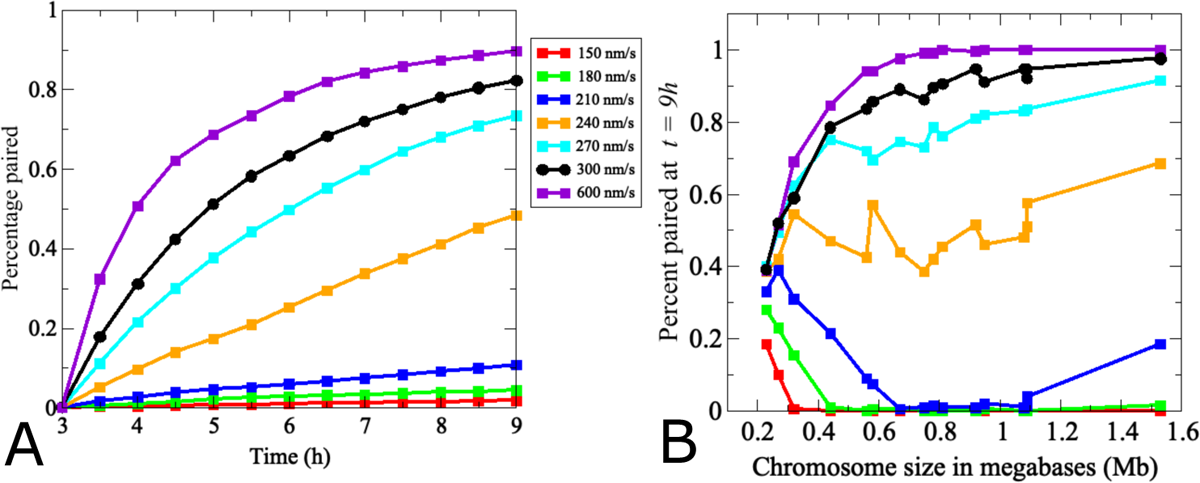
Modeling identifies a critical threshold of chromosome movement velocity for efficient homolog pairing. (A) Dots indicate the average pairing levels over time of all 32 true-sized chromosomes. Black indicates the velocity of chromosome movements in the wild-type model in Figs. 4 and 5 (300 nm/s). Chromosomes fail to pair at velocities around 150 nm/s. Increases in 30 nm/s increments improve pairing efficiencies at *t* = 9*h* 3-fold, with more modest gains above 240 nm/s where essentially all 16 homologs pair efficiently. For a more detailed analysis of pairing kinetics of (a) see Fig. S1 Fig. (B) Effects of movement velocity for different chromosome sizes. The graph shows pairing levels of chromosomes of increasing sizes at *t* = 9*h*, indicating that pairing efficiencies of larger chromosomes are more dramatically affected by changes in velocity than those of smaller chromosomes. Note that at higher velocities, chromosomes move further into the repulsive radii of their non-homologous partners and are therefore repelled faster, resulting in an accelerated homology search.

Thus, instead of a linear relationship between chromosome velocity and pairing, our model predicts a threshold effect where velocities at or above 240 nm/s dramatically improve pairing, a threshold that particularly impacts pairing of mid-sized and larger chromosomes. Together, these findings suggest that at velocities below 240 nm/s, contributions from random diffusion are more pronounced, interfering with directional chromosome movements. As random motion becomes dominant over directed movement, chromosomes explore less nuclear area, making homologous encounters less likely. Moreover, at lower translational velocities, repulsive interactions between non-homologous chromosomes are less frequent, diminishing effects of excluded regions defined by the presence of non-homologous chromosomes.

### Contributions of attractive and repulsive force strengths to homolog pairing

Next, we examined relative contributions of attractive and repulsive forces to pairing dynamics by appropriate parameter changes in Eqs. (1) to (3). The base model (Fig. 7A) comprising size adjusted chromosomes with equal strengths and reach of attractive as well as repulsive forces and movements at 300 nm/s was adjusted by setting either the attractive force strength *C_a_* = 0 (Fig.7B) or by inversely setting the repulsive force strength *C_r_* = 0 (Fig. 7C). These simulations indicate that neither attractive nor repulsive forces alone achieve pairing, at least on the time scale examined here, yet with some important differences. With only attractive forces, homologous chromosomes are drawn together only when they enter the local proximity of each other, and pairing proceeds exceedingly slowly, but with little or no effect of chromosome size (Fig.7C). In contrast, with repulsion only, the repulsive forces progressively drive homologs together via volume exclusion, with slower pairing kinetics affecting only smaller chromosomes (Fig.7B).

**Fig 7.**
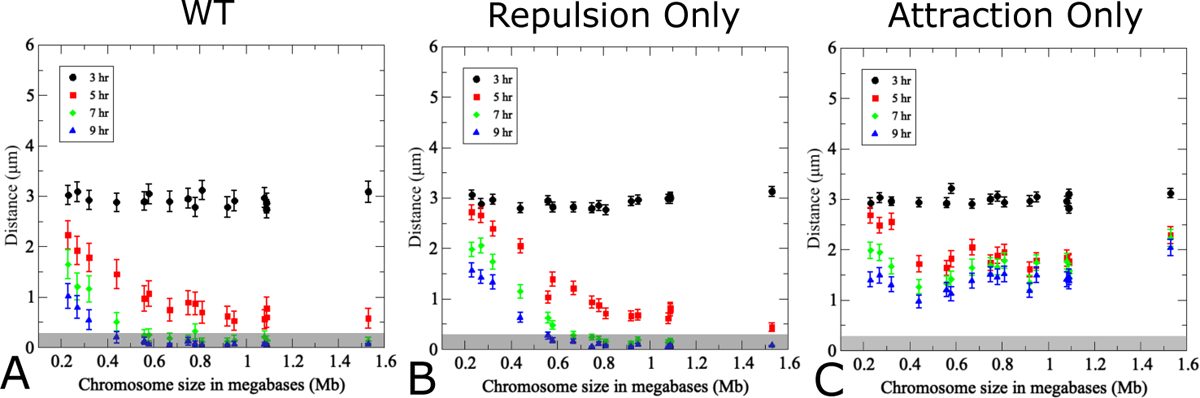
Contributions of attractive and repulsive forces on pairing efficiencies and kinetics. (A) Pairing wild-type model using true chromosome lengths and a standard translational movement velocity of 300 nm/s as primarily studied herein. In (B) the attractive strength *C_a_* = 0 in the WT model to highlight the effect of repulsive interactions alone. In (C) the reverse is true *C_r_* = 0 in the WT model to highlight the effect of attraction alone. (200 realizations, error bars indicate SD). The pairing distance is highlighted by a gray rectangle.

We also examined whether a dominant attractive force could result in homolog pairing by increasing the attractive force *C_a_* by an order of magnitude compared to the base model with an equivalent reduction of the repulsive force *C_r_*. Indeed, this parameter adjustment results in almost instantaneous pairing of all but the smallest chromosomes (see Supplemental Fig. S5 Fig). We note, however, that this scenario is somewhat unrealistic, as it is equivalent to a 3-fold increase in the effective radius, resulting in an effective radius of roughly 1200 nm for average-sized chromosomes, further corresponding to an average resection length of 2.4kb of fully extended single stranded DNA Supplemental Fig. S5 Fig). While symmetric attractive and repulsive forces more faithfully capture the dynamics observed in the reference experiment (Fig. 5), this permutation of the model demonstrates how systems with faster or slower pairing can be captured by adjusting the magnitude of both contributing interaction forces.

### Contributions of chromosome flexibility and orientation to homolog pairing

To better account for the rod-like shape of condensed prophase I chromosomes, we next wanted to extend the basic modeling framework to account for an approach more consistent with presumed polymer properties of chromosomes. Following an approach previously developed for linear active polymers [57], we modeled each chromosome as an active dumbbell where two beads, as defined by the base model, are connected by a bond force F*_b_* = ∇*U_b_*. Here the bond potential and resulting force are expressed as

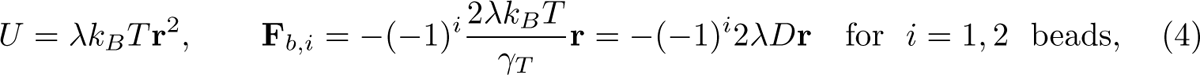

where *λ* is a Lagrange multiplier, *k_b_* is the Boltzmann constant, *T* is the temperature, *γ_T_* is a friction coefficient of translational motion and r^2^ = (r_2_ − r_1_)^2^ the bond vector whose average length enforces the chromosome size, (r^2^) = *ℓ*^2^. Following this approach we choose *k_b_T/γ_T_* = *D* the diffusion coefficient. This replaces the radial model above with an elastic dumbbell formed by two connected beads each with its own attractive and repulsive forces as above, but now explicitly adding the additional features of elasticity/flexibility and orientation of the chromosome (see Figure 8A). In the equations of motion the bond force term is added as an additional contribution to the motion of each chromosome, but only affects the relative position of each bead in the active dumbbell. In terms of attractive and repulsive interactions, each of the beads interacts with the two beads on each of the other chromosomes. This more faithfully represents the elongated physical structure of chromosomes.

**Fig 8.**
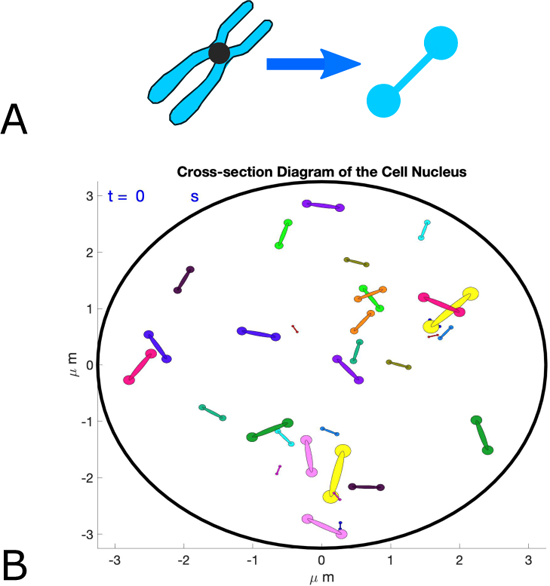
Polymer chain model for chromosomes incorporates flexibility and orientation. (A) Polymer chain model as an active dumbbell where each end is represented by the single bead model (1) with an additional term to ensure they stay together (4). (B) Representative still from a simulation of the active dumbbell movie, see Supplemental Movie S3 Video.

Simulations were carried out and described with a still of the nucleus and all 32 active dumbbell chromosomes (see Figure 9B and Supplemental Movie S2 Video). Even though pairing is completed somewhat faster in the model with dumbbells compared to those with circular chromosomes with a single center of mass, both models provide similar results. Thus, the computational complexity added by the active dumbbell approach does not significantly alter the results, at least with the parameters chosen here. However, the active dumbbell version of the simulation may prove useful in situations, e.g., when interactions with unusually large chromosomes are under investigation.

**Fig 9.**
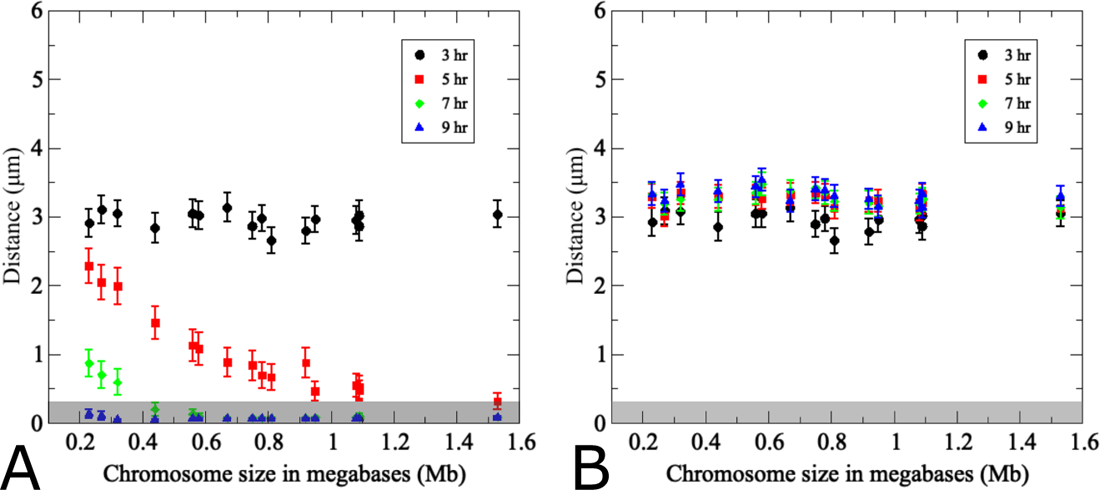
Polymer chain model as an active dumbbell consistent with simpler single bead model. (A) Average distances over time between homolog pairs where repulsive and attractive radii are adjusted proportionally to the sizes of actual yeast chromosomes. Note that the three smallest homolog pairs are markedly slower in achieving pairing than all other chromosomes. (B) Non-homologous chromosomes equidistant to each of the two homologs were identified at *t* = 3*h* and their distances were monitored throughout the simulation in the same set of model nuclei monitored in (A). (200 realizations, error bars indicate SD). The gray rectangle highlights pairing distances at or below 400 nm.

### Adaption of the pairing model to a mutant with reduced DSBs

Building on the expanded model that takes chromosome sizes and optimized velocities into account, we next investigated the case of a meiotic mutant for which experimental data are available [12]. Absence of meiotic DSBs, e.g., in a *spo11* null mutant, essentially abrogates homolog pairing, consistent with a central role of recombination intermediates in establishing and/or stabilizing homolog pairing [10, 16, 20]. A decrease of initiating DSBs to around 30% of wild type in a hypomorphic *spo11* mutant (*spo11-HA/spo11-HA-Y135F*; hereafter *spo11-HA/yf*) results in delayed, though largely efficient pairing [12]. Model parameters were adjusted to accommodate the fact that reduced DSB abundance would likely reduce both the cumulative attractive and repulsive forces exerted by these intermediates [9]. Furthermore, chromosome translation velocity was reduced consistent with experimental data that indicate a reduction of the average chromosome velocity in a *spo11* null mutant by 20% (to 110 nm/s from 140 nm/s in wild type; see Fig. 4A in [14]). Accordingly, for the *spo11* hypomorph, in one simulation we reduced the chromosome translation velocity from 300 nm/s to 230 nm/s.

Actual chromosome sizes were used for this simulation, yet results were plotted based on initial chromosome distances to facilitate comparison with the experimental data set in both wild-type *SPO11* (Fig. 10A) and *spo11* hypomorph (Fig. 10B,C). We considered two parameter sets to model the *spo11* hypomorph. In the first scenario, the movement speed was reduced to 77% of the wild type, and attractive strength wwas reduced 3-fold to represent the mutant strain’s decrease in DSBs available for the homology search (Fig. 10B). In the second scenario, we considered exclusively the DSB reduction in *spo11-HA/yf* by decreasing 3-fold the attractive as well as repulsive force strengths, but kept the chromosome movement speed at wild-type levels. For a specific realization of the homology search in mutant spo11, see Supplementary Movie S3 (https://zenodo.org/records/10246589). While reducing both movement speed and attractive forces delays pairing of widely separated homolog pairs indefinitely (see Fig. 10C), whereas all chromosome distances are affected more similarly when both attractive and repulsive contributions are reduced (Fig. 10B). Importantly, these responses demonstrate the versatility of our model to predict potential contributions of various parameters on pairing dynamics.

**Fig 10.**
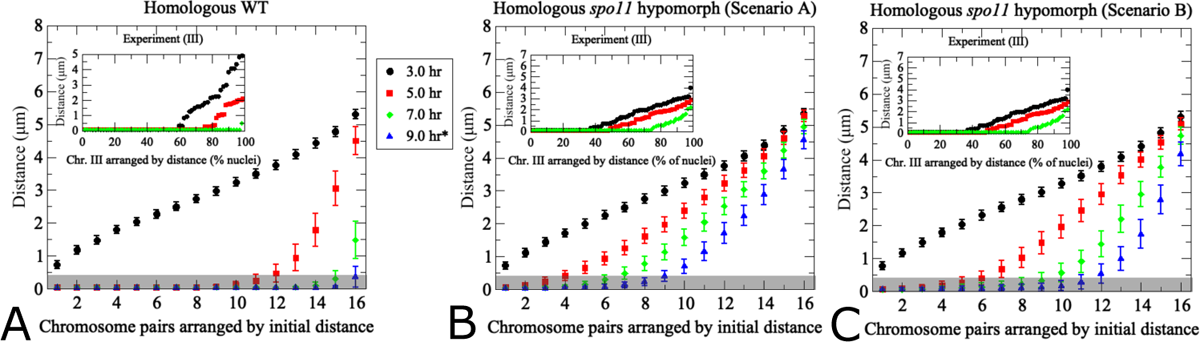
Modeling of pairing kinetics at reduced DSB abundance. (a) Wild-type model with true-sized chromosomes. Results differ from Fig. 4A) due to use of true sized rather than uniformly sized chromosomes. The inset shows experimental data for yeast chromosome III in hypomorphic *spo11-HA/yf*. Note the *x*-axis in the inset shows different nuclei derived from aliquots at the indicated time points, whereas the same cells were tracked through time during simulations. (b) Alternative model for hypomorphic *spo11* with 3-fold reductions of the wild-type levels of both attractive and repulsive force. For non-homologous chromosome distances see Fig. S3 Fig. (c) Model for *spo11* hypomorphic mutant with reduced movement speed and attractive force strength, with experimental observations for *spo11-HA/yf* shown in the inset. Translational movement velocity is reduced to 77% of wild type levels (230 nm/s), consistent with slower chromosome movements observed in *spo11* [14], and attractive force is reduced 3-fold representing decreased attractive forces exerted by fewer DSBs (see Table 1). The pairing distance is highlighted in gray at 400 nm. (200 realizations, error bars indicate SD).

## Discussion

How homologs pair during meiosis in the presence of an excess of non-homologous chromosomes is presently unknown. To explore contributions of the entire nuclear chromosome ensemble to the pairing process, we have developed an agent-based mathematical model derived from first principles that takes into account both attractive forces between homologs and dissociative forces between non-homologous chromosomes (see Methods section). In many organisms, homologs enter meiosis separated by distances that far exceed the reach of resected DSBs that could assess homology. This necessitates a process that ensures initial homolog co-localization potentially provided by chromosome movements together with non-homologous repulsive forces. Both attractive and repulsive forces exert their effects over the same short distances that are within the range of the single stranded region of a resected DSBs. Results from our simulations suggest that repulsive forces together with chromosome movements are a key determinant of bringing homologs into close vicinity. Repulsive interactions between non-homologous chromosomes create excluded regions within the nucleus driving homologs into close vicinity, thereby facilitating close-range attractive pairing interactions. Our simulations further demonstrate that repulsive forces are most effective when chromosome numbers rise above a certain threshold, likely by reducing the available area per nucleus. Attractive forces come into play once distances between homologs are sufficiently small.

A repulsive force that shortens the time spent in non-homologous interactions is a key feature of our mathematical pairing model. It represents molecular processes that dissociate non-homologous chromosomes from each other. While pronounced contributions of a dissociative/repulsive force to homolog pairing may appear counter-intuitive, several mutant phenotypes indicate the existence of molecular processes that contribute to the dissociation of non-homologous interactions which in the model are captured as a repulsive force.

First, when the heterodimeric Hop2/Mnd1 protein complex is defective, non-homologous chromosomes undergo stable synapsis in yeast, mammals and plants [36, 58–61]. The same protein complex also mediates homologous strand exchange [62], yet non-homologous synapsis is not a universal feature of mutants defective for strand exchange, indicating that elimination of non-homologous interactions represents a distinct function of the Hop2/Mnd1 complex [36] rather than being an indirect effect of defective strand exchange. Importantly, involvement of a single protein complex in homologous strand exchange and dissociation of non-homologous chromosome interactions is consistent with both forces acting upon the same molecular intermediate, most likely displacement loops between DNA segments with extensive or limited sequence similarity, respectively. Second, mutation of the Ph1 locus in allopolyploid wheat results in erroneous stabilization of interactions between non-homologous chromosomes, suggesting that Ph1 normally mediates disassociation of such interactions [63]. Third, the mismatch repair machinery disrupts recombination between DNA segments with limited sequence similarity via ejection of the invading strand [36, 39, 64]. In our model, this type of heteroduplex rejection is simulated by a repulsive force that minimizes the association time of non-homologous chromosome partners.

Repulsive interactions may also be the underlying cause for temporal separation of paired chromosomes observed in live-cell imaging studies [19]. Temporal separation of homologs has been interpreted and modeled as spatially restricted sub-diffusion involving fully paired homologs [19]. Movie animations of our model indicate that similar disruptions could arise due to collisions with non-homologous chromosomes during incipient pairing interactions of homolog pairs (e.g., see Supplemental Movie S1). Importantly, results from our model suggest that during this exploratory phase, homolog pairs are susceptible to becoming dislodged due to collisions with non-homologous chromosomes.

Earlier modeling approaches have focused on interactions between individual homologs which likely play an important role in completion of the pairing process, but they have not considered the effects of interactions between non-homologous chromosomes [19, 21–23]. The agent-based modeling framework developed here is new to chromosome dynamics but has been useful in modeling other multicomponent systems that involve attractive and repulsive forces, ranging from molecular to macroscopic components [21, 22, 27, 65–75].

Our model has identified unexpected threshold effects for several parameters where minor changes result in major nonlinear effects. Accordingly, for the current setting, homologous pairing levels and/or kinetics are disproportionately increased when the number of homolog pairs is increased from 2 to 4 (Fig. 3), when chromosome size exceeds 400 kbp (Fig. 5), and when chromosome velocity is increased from 210 nm/s to 240 nm/s (Fig. 6). Such discontinuities are likely related to a critical threshold of non-homologous chromosome encounters that needs to be crossed for homologous chromosomes to become confined to the same nuclear areas, thereby facilitating homolog encounters. Accordingly, when chromosomes are present in lower numbers, exhibit smaller sizes or fail to achieve unidirectional movement due to disturbance by Brownian motion, the frequency of non-homologous interactions is reduced and crowding of homologous chromosomes into the same nuclear area occurs at low frequencies.

Rapid chromosome movements during prophase of meiosis I have been observed in species from yeast to mouse [14, 15, 76]. Intriguingly, in different taxa, chromosome movements are mediated by different cytoskeleton components potentially resulting in a wide variation of movement speeds [29]. Our model indicates that chromosome movements must occur above a certain velocity threshold, likely determined by the number of chromosomes and the dimensions of the nucleus, providing a potential reason why chromosomes move at distinct speeds in different organisms. Moreover, our model predicts that the three smallest yeast chromosomes would be slower in completing homolog pairing (Fig. 5). Notably, the same chromosomes exhibit increased DSB and crossover frequencies [54, 55], features that may specifically compensate for size-related disadvantages in pairing.

A key feature of the current agent-based modeling is that all participating entities are included in the simulations, allowing for the ability to capture both typical behavior and deviations thereof. In contrast, experimental analysis of homolog pairing is limited by the availability of distinct tags for individual chromosomes. Accordingly, observations from a small number of homolog pairs have been extrapolated to the entire chromosome complement. Moreover, in cases where the pairing status is monitored in fixated cells, the progression of pairing must be inferred from different cells retrieved from the culture at different times. In contrast, our model captures pairing efficiencies and kinetics of all chromosomes in the same cell over time. This has already allowed us to predict different sensitivities to translation velocity thresholds of small, medium-sized, and very large chromosomes that would have eluded a population-based approach (see Fig. 6B).

One of the key features of our modeling framework is that it is easy to further explore the parameter space. Our model is readily scaled to different biological settings and may provide predictions for a multitude of cellular scenarios, laying the groundwork for experimental studies. For example, meiosis in the Indian muntjac involves only three very large homolog pairs, whereas in some insects and plants between 300 to 600 homolog pairs need to complete pairing [77, 78]. Such complexities are inaccessible to current experimentation but become analyzable by the current model. In future work, our model could be used to explore, e.g., how pairing dynamics are affected by different nucleus sizes and shapes, or by a nucleus represented by a 3-dimensional volume rather than a 2 dimensional surface area. As an example, we have already started exploring the effect of nucleus size on pairing dynamics, indicating that a smaller nuclear area accelerates pairing likely because each chromosome interacts with several non-homologous neighbors at the same time.

Conversely, pairing becomes inefficient when the nuclear area is increased above a certain size (e.g., Fig. S4 Fig). Other extensions of our model might include chromosome-size dependent velocities, and temporary changes in effective nuclear volume, as in the case of directed chromosome movements during the horsetail stage in *S. pombe* where the entire chromosome complement temporarily becomes confined to small regions within the nuclear volume.

In summary, the modeling approach developed here suggests that homolog pairing is achieved by two mechanistically distinct, yet temporally coinciding processes: Homologs become confined to a nuclear area due to the dissociation of interactions with the entire non-homologous chromosome set achieved via a repulsive force.

Confinement to smaller areas enables homologs to assess similarities modeled as an attractive force. Importantly, both types of interactions involve close-range physical DNA interactions. Our model makes specific predictions about contributions of chromosome dimensions and movement velocity in combination with chromosome numbers that may further be affected by specific nuclear dimensions. Chromosome number, nuclear dimensions and movement speeds vary widely among different organisms and may affect pairing requirements. All three parameters are accessible to experimental manipulation [78], rendering predictions by our model testable in appropriate experiments.

## Supporting information

### Effect of velocity on pairing kinetics

Similar to Fig. 3, we consider the effect of movement velocity on pairing kinetics. In particular, we observe that pairing kinetics are accelerated as velocity increases. Fig. S1 FigA is a reproduction of Fig. 6A) showing how movement velocity changes the homolog pairing efficiency. Fig. S1 FigB, rescales each curve in (A) by the maximum pairing efficiency. The model predicts that the benefits associated with increased velocity appear to saturate past 300 nm/s. To investigate this trend’s direct effect on pairing kinetics, we compute the distances between homologs as a function of chromosome movement speed. The results are summarized in Fig. S2 FigA-D. Accordingly, smaller compared to larger chromosomes require lower movement velocities for timely pairing completion.

S1 Fig. Effects of chromosome velocity on pairing kinetics. (A) Reproduction of Fig. 6a, each dot indicates the average pairing levels of all 32 chromosomes of true sizes at the indicated time points. Black (300 nm/s) indicates pairing levels at the velocity of chromosome movements in the wild type model in Figs. 3-5 Chromosomes fail to pair at velocities around 150 nm/s, likely due to the effect of thermal noise. Increases in 30 nm/s increments increases pairing efficiencies at *t* = 9*h* 3-fold, with more modest gains above 240 nm/s where essentially all 16 homologs pair efficiently. (B) Results from Fig. S1A were normalized by maximum pairing levels, as described for Fig. 3B. With increased movement velocities, 50% pairing levels are achieved at progressively earlier time points.

S2 Fig. Effects of movement velocity on homologous pairing kinetics. All results for 16 homolog pairs using true chromosome lengths. Chromosomes are arranged according to their initial distance to facilitate comparison with the experimental data set. Chromosome movement velocity is (A) 180 nm/s, (B) 210 nm/s, (C) 240 nm/s, and (D) 270 nm/s. Note the sharp transition between 210 nm/s and 240 nm/s, as also shown in Fig. 6A. The pairing distance is highlighted in gray at 400 nm. (200 realizations, error bars indicate SD).

### Non-homologous chromosome distances in wild type compared to *spo11* hypomorph

The complementary plot to Fig. 7 shows the distribution of non-homologous pairing distances as a function of time. Fig. S3 Fig provides a direct comparison between non-homologous chromosome distances in wild type (A) and *spo11* hypomorph (B) where true chromosome lengths are used. Numerically, we tracked the distance between a given homolog and selected the non-homologous chromosome in the same nucleus with the closest initial distance to its homologous pair. The simulation then tracks the dynamics of both throughout time and we plot the distance between a given chromosome and the previously identified non-homologous chromosome. The results indicate that there is no bias in time for a chromosome with any non-homologous chromosome in the nucleus. Also, the results indicate that the mutation does not have a strong effect on non-homologous interactions as it only modifies their strength, but not their qualitative behavior of creating excluded regions. One difference as noted in Fig. 7 is that while both the wild type and mutant produce excluded regions, the strength of the repulsion in the wild type makes these regions more severe and drives homologous pairing to occur much faster. This validates the model in that the only different interaction occurs with a given chromosome’s homologous mate (compare homologous results in Fig. 7 with non-homologous results in Fig. S3 Fig).

S3 Fig. Non-homologous chromosome distances for wild type and *spo11* hypomorph, with experimental observations in the insets. (A) Reproduction of Fig. 4B for non-homologous distances in the wild type simulations. (B) Non-homologous pairs of *spo11* mutants where one of the two homolog partners is matched with a non-homologous chromosome that exhibits an optimally matched initial distance with its cognate homolog partner at *t* = 3*h*. Inset in (B) shows experimentally determined distances between non-homologous GFP tagged chromosomes II and III. For details on the experimental conditions see [12]. The pairing distance is highlighted in gray at 400 nm. *Note the experimental data do not include information for *t* = 9*h*. (200 realizations, error bars indicate SD).

### Effect of nucleus size on chromosome kinetics

S4 Fig. Effects of nucleus size on pairing kinetics and efficiency. Average distances over time between homolog pairs to the sizes of actual yeast chromosomes. Here we compare the dynamics as the nucleus size is varied. Observe that pairing occurs much less frequently as the nucleus radius increases. This indicates that confinement and the repulsive non-homologous interactions play a critical role in the kinetics. In a high-density environment like the small nucleus the repulsive interactions have an even larger effect because each chromosome is interacting with many non-homologous neighbors in close proximity. This drives the pairing of homologs by quickly filling the nucleus space with excluded regions due to repulsion. Results over 200 realizations.

### Effect of order of magnitude increase/decrease of interaction forces

Here we consider scenarios where either the attractive force strength *C_a_* dominates 100-fold over the repulsive force strength *C_r_* (Fig. S5 FigB) or inversely, the repulsive force strength dominates over the attractive force strength (Fig. S5 FigC). Moreover, with the attractive force dominating, pairing of mid-sized and longer chromosomes occurs essentially instantaneously, whereas the process is drawn out over a longer time scale when repulsive forces dominate.

S5 Fig. Contributions of attractive and repulsive forces on pairing efficiencies and kinetics. (A) Pairing wild-type model using true chromosome lengths and a standard translational movement velocity of 300 nm/s as primarily studied herein. In (B) the repulsive strength is increased by an order of magnitude *C_r_* = 0.05 and the attractive strength is decreased by an order of magnitude *C_a_* = 0.0005. In (C) the reverse is true *C_r_* = 0.0005 and *C_a_* = 0.05 (200 realizations, error bars indicate SD). The pairing distance is highlighted by a gray rectangle.

### Supplemental movie files

Movie files are available at a dedicated online Zenodo Repository: https://zenodo.org/records/10246589

S1 Video. WT Chromosome Trajectories during Prophase I. The movie shows one realization of the agent-based model. The simulation movie covers the homology search process from *t* = 3*h* to *t* = 9*h*. Matching colors correspond to homologous pairs. True chromosome lengths are incorporated and scale the relevant interaction radii. The radius represents the attractive and non-homologous repulsive region.

S2 Video. WT Chromosome Trajectories during Prophase I with active dumbbell model. The movie shows one realization of the agent-based model active dumbbell model which is closer to modeling a chromosome as a polymer. The simulation movie covers the homology search process from *t* = 3*h* to *t* = 9*h*. Matching colors correspond to homologous pairs. True chromosome lengths are incorporated and scale the relevant interaction radii, but are allowed to change in time as the two beads expand and contract. The radius represents the attractive and non-homologous repulsive region.

S3 Video. spo11 hypomorph (30% WT DSB levels) Chromosome Trajectories during Prophase I (parameters from Fig. 7B). The movie depicts one realization of the agent-based model for the *spo11* hypomorphic mutant. The simulation movie covers the homology search process from *t* = 3*h* to *t* = 9*h* where mutant *spo11* is associated with a weaker attractive and repulsive force (e.g., reduction to 77% of WT values). True chromosome lengths are incorporated and scale the relevant interaction radii. Matching colors correspond to homologous pairs. The radii represent the homologous attractive and the non-homologous repulsive region. Note that the reduction in interaction strength delays homologous pairing consistent with experimental observations in [12].

## Acknowledgments

The work of G.V.B. was supported by the National Institute of General Medical Sciences of the National Institutes of Health under Award Number R01GM141698. The content is solely the responsibility of the authors and does not necessarily represent the official views of the National Institutes of Health. The funders had no role in study design, data collection and analysis, decision to publish, or preparation of the manuscript. The authors thank Sebastian Sensale Rodriguez for useful discussions. AC and SR developed the mathematical model and carried out numerical simulations, emerging from discussion with GVB. AC, GVB, and SR analyzed the results and wrote the manuscript.

